# Antigen specificity of clonally-enriched CD8+ T cells in multiple sclerosis

**DOI:** 10.1101/2024.09.07.611010

**Authors:** Kristen Mittl, Fumie Hayashi, Ravi Dandekar, Ryan D. Schubert, Josiah Gerdts, Lindsay Oshiro, Rita Loudermilk, Ariele Greenfield, Danillo G. Augusto, Akshaya Ramesh, Edwina Tran, Kaniskha Koshal, Kerry Kizer, Joanna Dreux, Alaina Cagalingan, Florian Schustek, Lena Flood, Tamson Moore, Lisa L. Kirkemo, Tiffany Cooper, Meagan Harms, Refujia Gomez, University of California, San Francisco MS-EPIC Team, Leah Sibener, Bruce A. C. Cree, Stephen L. Hauser, Jill A. Hollenbach, Marvin Gee, Michael R. Wilson, Scott S. Zamvil, Joseph J. Sabatino

## Abstract

CD8+ T cells are the dominant lymphocyte population in multiple sclerosis (MS) lesions where they are highly clonally expanded. The clonal identity, function, and antigen specificity of CD8+ T cells in MS are not well understood. Here we report a comprehensive single-cell RNA-seq and T cell receptor (TCR)-seq analysis of the cerebrospinal fluid (CSF) and blood from a cohort of treatment-naïve MS patients and control participants. A small subset of highly expanded and activated CD8+ T cells were enriched in the CSF in MS that displayed high activation, cytotoxicity and tissue-homing transcriptional profiles. Using a combination of unbiased and targeted antigen discovery approaches, MS-derived CD8+ T cell clonotypes recognizing Epstein-Barr virus (EBV) antigens and multiple novel mimotopes were identified. These findings shed vital insight into the role of CD8+ T cells in MS and pave the way towards disease biomarkers and therapeutic targets.

**Graphical Summary.**
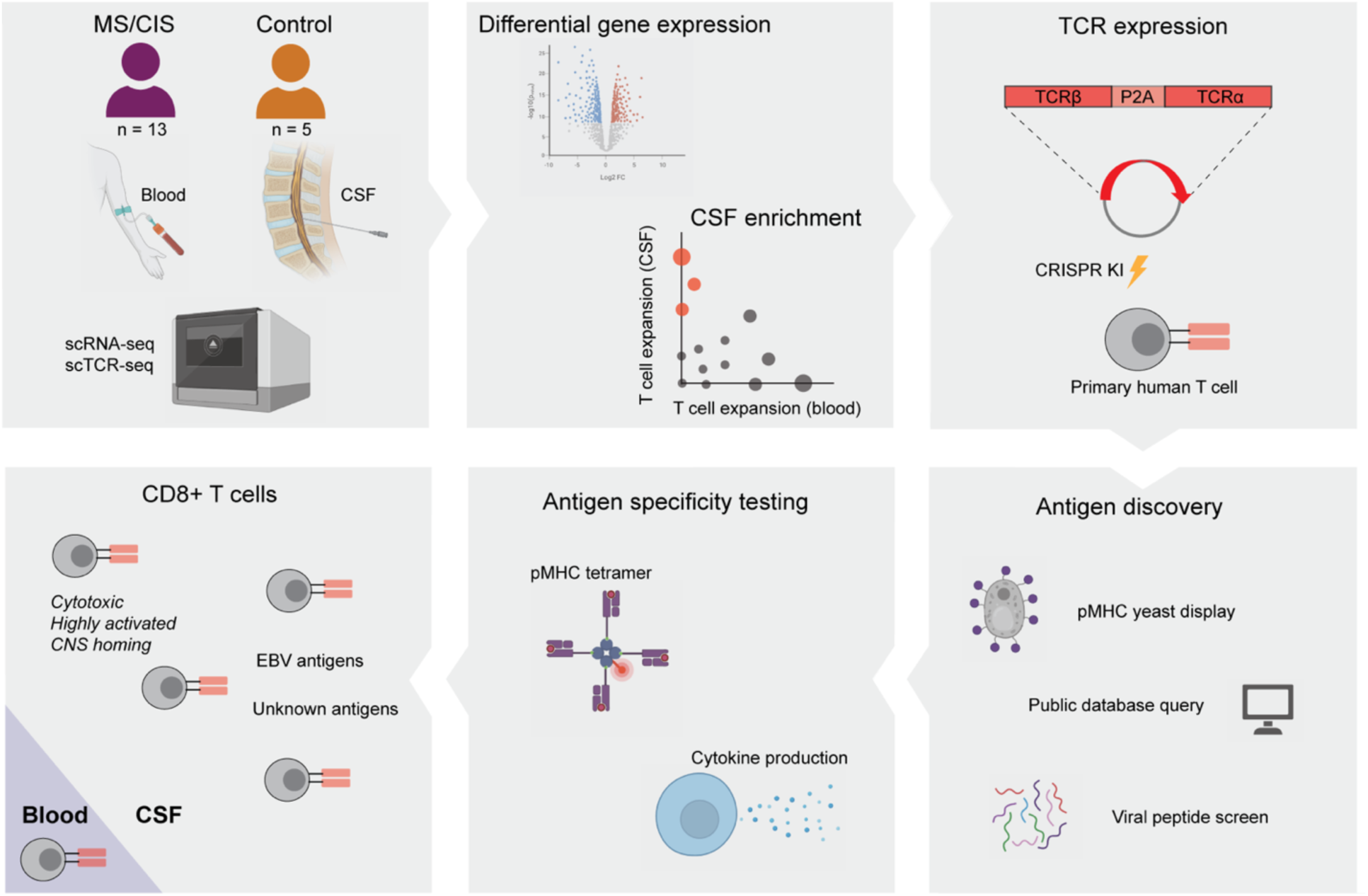
Created in BioRender. Sabatino, J. (2024) BioRender.com/e66l598

## INTRODUCTION

Multiple sclerosis is an inflammatory demyelinating condition of the central nervous system (CNS) characterized by significant T cell involvement^1^. Both CD4+ and CD8+ T cells are found within the perivascular spaces as well as in the parenchyma of MS lesions^2–4^. CD8+ T cells are enriched and clonally expanded in MS lesions relative to CD4+ T cells^5–8^, suggesting local antigen-driven expansion. Specific MHC I alleles also alter MS susceptibility^9^, providing additional support for a critical role of cytotoxic CD8+ T cells in MS biology. The goal of this study was to identify T cells, particularly CD8+ T cells, that are uniquely expanded within the central nervous system (CNS) and determine their phenotypic characteristics and antigen specificity.

Acquiring CNS tissue to analyze the infiltrating T cell response in MS is invasive and rarely performed early in the disease course. Cerebrospinal fluid (CSF) is a transit site of lymphocytes entering the CNS^10,11^ and provides a window into the immune responses within the CNS. CSF repertoire studies have indicated high clonal overlap with expanded T cell populations within MS lesions^7,12,13^, therefore we performed T cell clonal analysis in the CSF compared to the peripheral blood to characterize the immune response within the CNS.

Single-cell sequencing technologies provide powerful opportunities for deep phenotyping and clonal characterization of T cells in numerous diseases, including MS^14–17^, yet detailed analysis of CSF-expanded T cells and their antigen specificity in MS are currently lacking. In this study, single-cell RNA-sequencing (scRNA-seq) paired with single-cell T cell receptor-sequencing (scTCR-seq) was employed from the CSF and blood from individuals with early untreated MS and control participants. After identifying a subset of CSF-expanded CD8+ T cells in MS patients, their antigen specificity was investigated using a combination of unbiased and targeted antigen discovery methodologies.

## RESULTS

### Study participants

18 individuals were enrolled in the study: 11 relapsing-remitting MS (RR-MS), 2 clinically isolated syndrome (CIS), 2 other neuroinflammatory disorders (OND: neurosarcoidosis and intermediate uveitis), and 3 healthy controls (HC). Demographics are shown for each cohort in **Table 1** and each individual in **Table S1.** All of the MS/CIS patients were treatment-naïve (i.e. no prior history of immunomodulatory or immunosuppressive therapies) at the time of sample collection, but one OND patient was on immunotherapy (TNFα inhibitor).

**Table 1.** Overview of participant characteristics. OCB+ refers to positive oligoclonal band status. Percent active MRI refers to contrast enhancement on MRI within 30 days of blood/CSF analysis.

| **Disease** | **N** | **Gender (F/M)** | **Mean Age** | **% Untreated** | **% OCB+** | **% active MRI** |
| --- | --- | --- | --- | --- | --- | --- |
| RR-MS | 11 | 7/4 | 41 | 100% | 82% | 64% |
| CIS | 2 | 2/0 | 44 | 100% | 0% | 0% |
| OND | 2 | 2/0 | 35 | 50% | 100% | Not tested |
| HC | 3 | 1/2 | 34 | N/A | 0% | Not tested |

### Identification of T cell subsets in the blood and CSF by single-cell sequencing

Paired peripheral blood and CSF were collected on the same day for each study participant. Freshly acquired samples comprised of all unseparated cell subsets underwent paired scRNA-seq and scTCR-seq using the 10X Genomics 5’ library preparation kits to permit combined single-cell transcriptional phenotyping and TCR clonal analysis. The scRNA-seq data for all participants in this study was previously published^18^. All major immune cell subsets were readily identified from the scRNA-seq data, with T cell clusters comprising the largest fractions (**Fig. 1A**). To characterize conventional TCRαβ T cells, all subsequent analyses focused on only those T cells with paired scRNA-seq and scTCR-seq data (**Fig. 1B**). In total, 48,468 individual T cells were identified from the blood and CSF across all participants (**Fig. 1C**). As expected, TCR-associated genes (*CD3E, CD3D*) were highly upregulated with absent expression of non-T cell-associated genes (e.g. *CD19*) (**Fig. S1**). 33,349 CD4+ and 15,119 CD8+ T cells expressing paired TCRαβ genes were identified (as described in Materials and Methods) for analysis from the combination of blood and CSF of all 18 participants (**Table S2**).

**Fig. 1.**
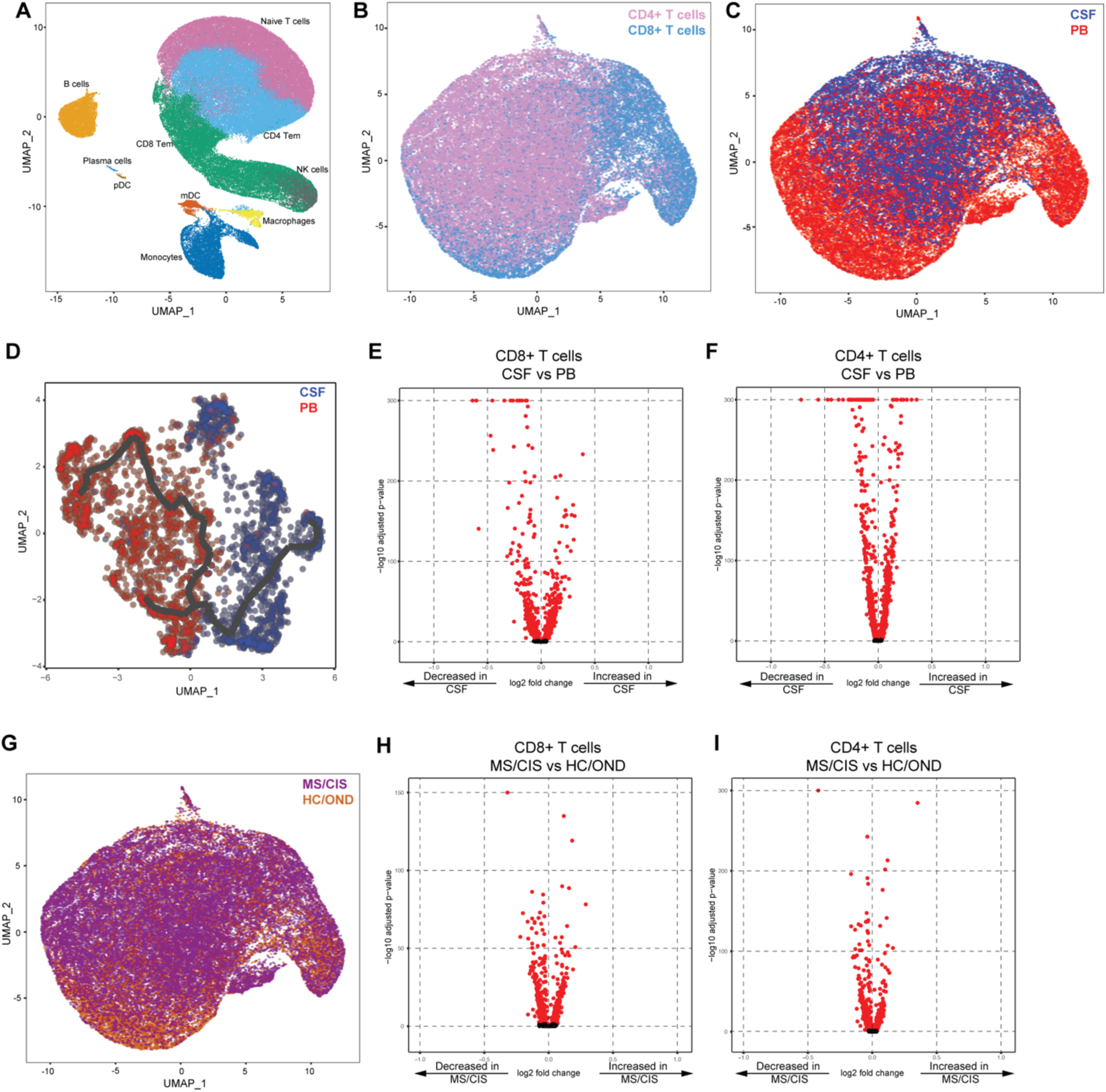
T cell single cell sequencing analysis in blood and CSF. Major immune cell subsets from combined blood and CSF of all patients were identified by scRNA-seq (**A**). T cells were defined after integration of scRNA-seq and scTCR-seq data, allowing segregation of T cells by CD4/CD8 status (**B**), compartment (CSF) (**C**), and disease status (**G**). Pseudotime trajectory analysis of CSF and PB is shown in **D**. Volcano plot analysis of differential gene expression between the CSF and PB for CD8+ T cells (**E**) and CD4+ T cells (**F**) and between MS/CIS and HC/OND for CD8+ T cells (**H**) and CD4+ T cells (**I**). Genes with adjusted p-values < 0.05 are indicated in red. Abbreviations: PB = peripheral blood; CSF = cerebrospinal fluid; MS = multiple sclerosis; CIS = clinically isolated syndrome; HC = healthy control; OND = other neuroinflammatory disorder.

Pseudotime analysis revealed distinct populations of T cells largely segregated based on compartment (i.e. CSF or blood), highlighting the distinct transcriptional signatures associated with different anatomic sites (**Fig. 1D**). Both CD8+ and CD4+ T cells were distinct between the peripheral blood and CSF (**Fig. 1E-F**). For instance, CD8+ T cells (**Fig. 1E, Table S3**) in the CSF displayed significantly increased expression of various genes relative to the peripheral blood, including genes associated with migration and trafficking (*CXCR3*, *CXCR4*, *CCL4*, *ITGB1, ITGA4*), signaling and activation (*CD2*, *FYN*, *DUSP2*), and cytotoxicity (*GZMK*, *GZMA*). In contrast, peripheral blood CD8+ T cells expressed significantly higher levels of *FOS*, *JUN*, *DUSP1*, and *GADD45B* amongst other genes compared to the CSF, indicating an alternate activation state. Within the CSF, CD4+ T cells (**Table S3**) showed significantly increased expression of genes similar to their CD8+ T cell counterparts (*ITGB1*, *ITGA4*, *CXCR3*, *GZMA*, *GZMK*, *CD2*) as well as distinct genes (*JUN, FOS, DUSP1, CCR7, HCST*).

Given the disproportionate number of participants in different disease categories (**Table 1**), we grouped MS and CIS patients together (n = 13) and performed differential gene expression against OND and HC combined as a comparison non-MS group (n = 5) (**Fig. 1G**). Within CD8+ T cells combined from the peripheral blood and CSF, various genes were differentially expressed between MS/CIS and HC/OND participants (**Fig. 1H**). In particular, genes associated with tissue trafficking (*CXCR4*, *CCL5*, *KLF2*, *ITGA4*, *ITGB1*, *CD69*) and cytotoxicity (*GZMK*, *KLRG1*, *GZMA*) were upregulated in MS/CIS (**Table S4**). In contrast, genes associated with naïve status (*CCR7*, *SELL*, *LEF1*) and TCR signaling (*CD8B*, *CD3D*, *CD3E*, *LCK*, *ZAP70*, *LAT*) were downregulated relative to HC/OND participants. A similar profile was observed amongst CD4+ T cells from blood and CSF of MS/CIS patients, including increased expression of genes related to tissue migration (*ITGB1*, *CD69*, *ITGA4*, *CXCR3*, *CXCR4*) and cytokine secretion (*IL32*, *GZMK*) and reduced naïve status (*CCR7*, *LEF1*, *SELL*, *TCF7*) and TCR signaling (*LCK*) (**Fig. 1I**, **Table S4**). Overall, these data suggest that both CD8+ and CD4+ T cells in MS/CIS patients are more activated with increased effector functions and tissue homing capacity than HC/OND participants, consistent with other studies^15,16,19^.

### T cell clonal analysis

The clonal repertoire of T cell subsets across compartments (i.e. peripheral blood versus CSF) and across disease states (i.e. MS/CIS versus HC/OND participants) was compared. T cell clonotypes were defined as T cells sharing identical V and J genes and CDR3 amino acid sequences for paired TCRαβ sequences similar to prior studies^14,20^. A total of 31,756 unique CD4+ T cell clonotypes and 10,825 unique CD8+ T cell clonotypes were identified from all individuals (**Table S2**). CD4+ and CD8+ T cell diversities (measured by Shannon entropy) were significantly higher in MS/CIS compared to HC/OND in the peripheral blood and CSF (**Fig. S2**). The diversity of CD4+ T cells was significantly higher in blood compared to CSF, but not for CD8+ T cells (**Fig. S2**). These findings suggest a diverse array of T cell clonotypes may be preferentially recruited in both the blood and CSF of MS/CIS patients.

T cell clonal expansion is a hallmark of prior antigen encounter, therefore our analysis focused on T cell clonality in the CSF. T cells were divided into three different categories of clonal expansion: unexpanded (single-cell of a given clonotype), moderately expanded (> 1 cell, but < 0.75% of an individual’s CSF T cell repertoire), or highly expanded (≥ 0.75% of an individual’s CSF T cell repertoire). We chose 0.75% as a cut-off for highly expanded clonotypes as it represented a relatively high threshold based on the clonal expansion observed in the CSF of our patient cohorts as well as others^14,21^. While small fractions of clonally expanded CD4+ T cells were observed in the CSF, much larger populations of highly and moderately expanded CD8+ T cells were observed in similar proportions in MS/CIS and HC/OND participants (**Fig. 2A**).

**Fig. 2.**
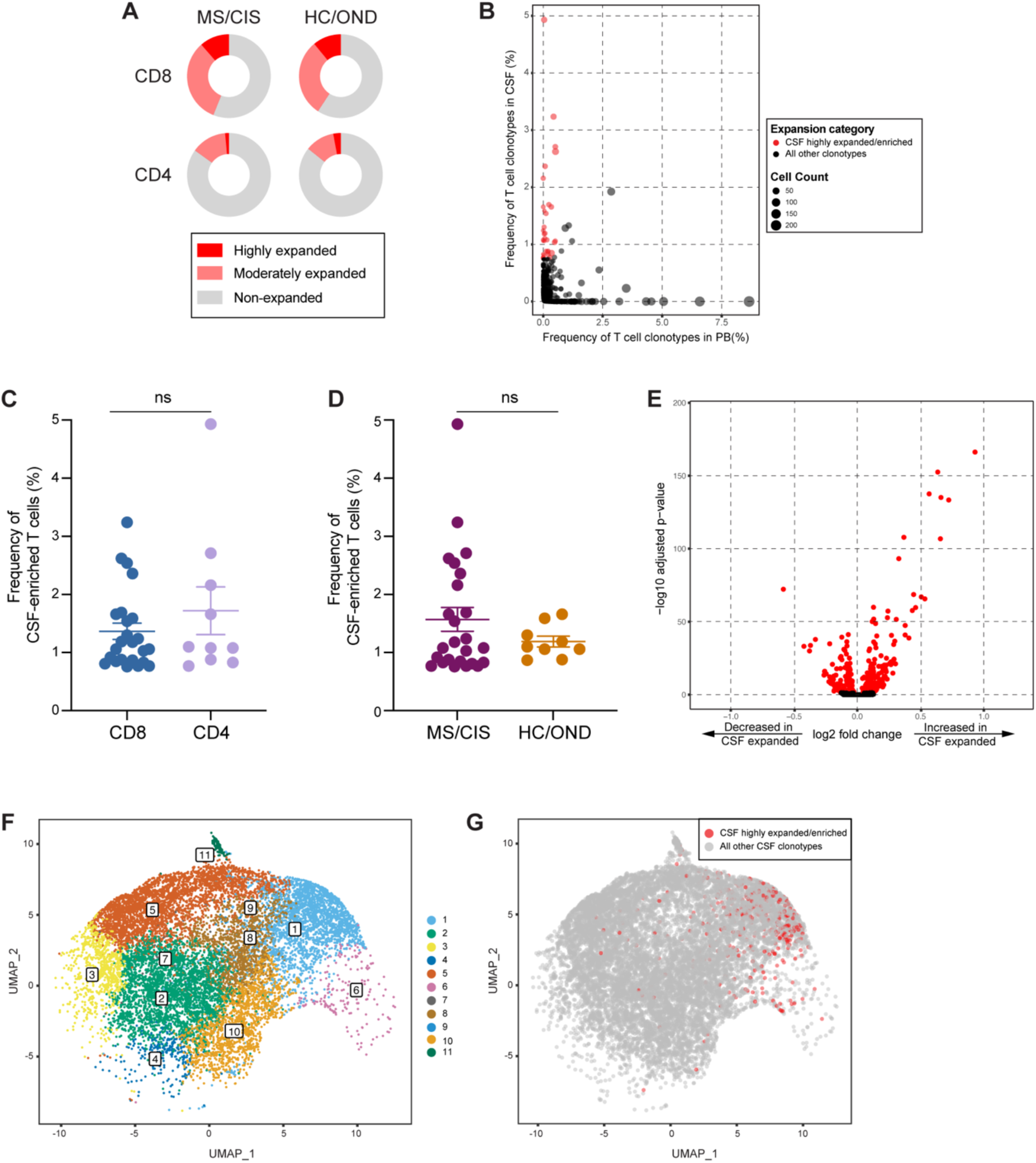
T cell clonal expansion in CSF. CD8+ and CD4+ T cell clonal expansion was compared between MS/CIS and HC/OND subjects (**A**). Non-expanded are T cell clonotypes that were present at singletons, moderately expanded clonotypes were more than one but less than 0.75% of the CSF repertoire and highly expanded T cell clonotypes comprised at least 0.75% of the CSF repertoire in a given individual. Clonal frequency of all T cell clonotypes in the CSF and blood that were highly expanded T cells and enriched at least 2-fold more frequently than the blood of the same individual are highlighted in red (**B**). The frequencies of highly expanded and enriched T cells is shown for CD8/CD4 status (**C**) and disease status (**D**). Volcano plot analysis of differential gene expression between highly expanded and non-expanded T cells in the CSF where genes with adjusted p-values < 0.05 are indicated in red (**E**). Unbiased clustering of all CSF T cells (**F**) overlaid with highly expanded/enriched T cells (**G**).

### Selective enrichment of highly expanded T cell clonotypes in the CSF

To delineate between T cells expanded comparably in the blood and CSF versus those preferentially expanded in the CSF, the abundance of all T cell clonotypes in the blood and CSF was compared in all individuals. The overwhelming majority of T cell clonotypes were detected in the blood or CSF only, whereas only ∼1.5% of all clonotypes were found in both compartments (**Fig. 2B**). We postulated that highly expanded T cell clonotypes (i.e. a CSF frequency ≥ 0.75%) that were enriched in the CSF relative to the peripheral blood were likely to be responsive to local antigens in the CNS. Enriched CSF-expanded T cell clonotypes were defined as those with a CSF frequency at least two-fold greater than the peripheral blood frequency from the same individual. This yielded 33 highly CSF-enriched and expanded T cell clonotypes varying from approximately 2-fold to more than 100-fold higher frequencies in the CSF relative to peripheral blood (**Fig. 2B**, **Table S5**). More than 70% of the highly expanded and CSF-enriched T cell clonotypes in the CSF were CD8+ T cells. The frequencies of highly expanded CSF-enriched CD8+ and CD4+ T cells were similar, ranging from 0.76-4.9% of an individual’s entire CSF repertoire (**Fig. 2C)**. Although there were no statistically significant differences in the mean frequencies of highly expanded CSF-enriched T cell clonotypes between MS/CIS and HC/OND, only MS/CIS participants had CSF-enriched T cells with frequencies greater than 2% (**Fig. 2D**). One MS patient, MS6, had 11 highly enriched T cell clonotypes, the majority of which were CD8+ T cells, and encompassing nearly 20% of their CSF repertoire (**Table S5**).

### Single-cell transcriptomics of CSF-expanded T cells

Highly expanded and unexpanded T cells in the CSF were compared by scRNA-seq analysis. Substantial differential gene expression changes were observed in highly expanded T cells in comparison to their unexpanded counterparts (**Fig. 2E**, **Table S6**). In particular, genes associated with cytotoxic CD8+ T cell function (*CD8A*, *CD8B*, *NKG7*, *KLRD1*, *GZMA*, *GZMH*, *GZMM*, *GZMK*, *EOMES*) and chemotaxis (*CCL5*, *CCL4*) were significantly increased in highly expanded T cells, whereas genes associated with naïve status were significantly reduced (*IL7R*, *LTB*, *LDHB*). Targeted gene expression analysis revealed increased expression of genes associated with effector/memory differentiation (*EOMES*, *KLRG1*, *CCL5*, *CD27*), tissue homing (*CXCR3*, *CCR5*), resident memory status (*CD69*, *IGTAE*), as well as inhibitory genes associated with chronic antigen exposure (*HOPX*, *TIGIT*, *DUSP2*, *PDCD1*, *LAG3*) in highly CSF-expanded CD8+ and CD4+ T cells (**Fig. S3**). In contrast, CSF-unexpanded T cells expressed higher levels of genes associated with naive/non-activation (*SELL*, *CCR7*, *IL7R*, *TCF7*, *LEF1*) as well as the integrin gene *ITGB1*. To further characterize CSF-enriched and expanded T cell clonotypes, the 33 T cell clonotypes were overlaid with 11 distinct CSF T cell clusters (**Fig. 2F-G**). The overwhelming majority of the enriched and expanded clonotypes were found in cluster 1, which was defined by a significantly increased expression with a number of genes associated with cytotoxic effector CD8+ T cells, including *CD8A*, *CD8B*, *PLEK*, *DUSP2*, *EOMES*, *GZMK*, *GZMA*, *GZMH*, *PRF1*, *NKG7*, *CCL5*, and *CCL4* (**Table S7**).

### Clonal relationships of expanded CSF-enriched T cells

Nearly all CSF T cell clonotypes across all individuals were unique. Only 21 identical TCRs (i.e. same V and J genes and CDR3 amino acid sequences for the paired α and β chains) were found between the peripheral blood of different individuals and another 3 that were identical between the blood and CSF of different individuals, irrespective of disease status (**Fig. 3A**). To further assess clonal relationships, GLIPH2 was employed, an algorithm to help identify TCRs with potentially shared specificity based on sequence similarity within the CDR3β region^22^. GLIPH2 was performed on all CSF T cell clonotypes, and the output was then queried against the 33 CSF high enriched CDR3 sequences. Using this approach, 19 clonally related networks comprised of 44 total clonotypes were identified (**Fig. 3B; Table S8**). The majority of networks comprised two related clonotypes; two networks were comprised of five clonotypes each. Almost all networks consisted of clonotypes from the same individual and were identified primarily amongst MS and CIS participants (**Fig. 3B**). Nearly all of the clonally-related T cells were CD8+ T cells (**Table S8**), suggesting potential shared antigen specificity.

**Fig. 3.**
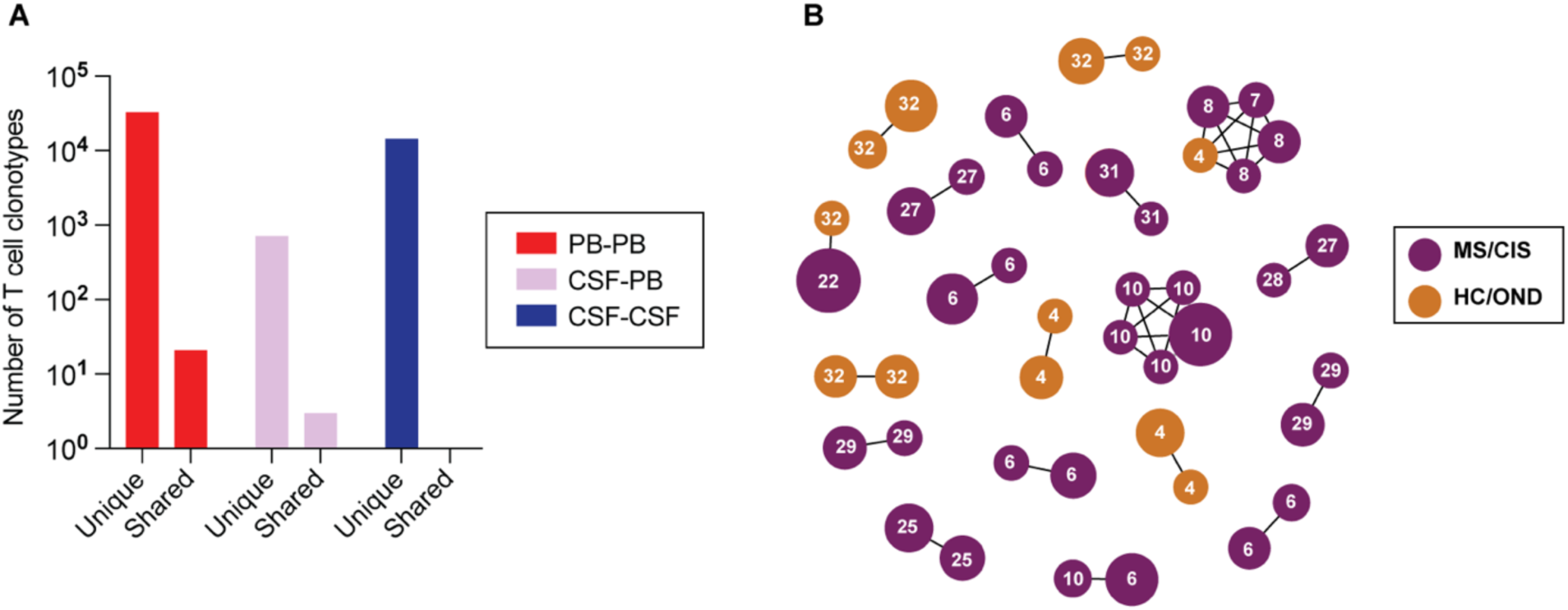
T cell clonal relationships. The number of unique or shared clonotypes between different compartments across different individuals is shown (**A**). GLIPH2 analysis of highly expanded and enriched T cell clonotypes in the CSF compared to all other CSF T cell clonotypes (**B**). Clonal size is indicated by node size and clonally related populations are connected by lines. The number in each clonotype refers to the subject ID.

### Antigen discovery of highly expanded CSF-enriched CD8+ T cells

Antigen discovery efforts focused on the 23 CD8+ T cell clonotypes since they comprised more than 70% of the expanded CSF-enriched T cells. Several different strategies were undertaken to explore the potential antigenic targets of these CD8+ T cell clonotypes (**Fig. 4A**). An unbiased antigen discovery approach was first employed using a peptide:MHC (pMHC) yeast display library in which ∼10^8^-10^9^ random peptides are displayed on a given MHC allele for probing recognition against individual TCRs^23^. 15 of 23 CD8+ TCRs were successfully expressed and tested against specific MHC I allele libraries based on library availability and the alleles of the participants from which the TCRs were derived (**Table S9**). Four TCRs (3 MS/CIS, 1 HC) demonstrated substantial enrichment of specific peptides from three different MHC I libraries (**Fig. 4B**).

**Fig. 4.**
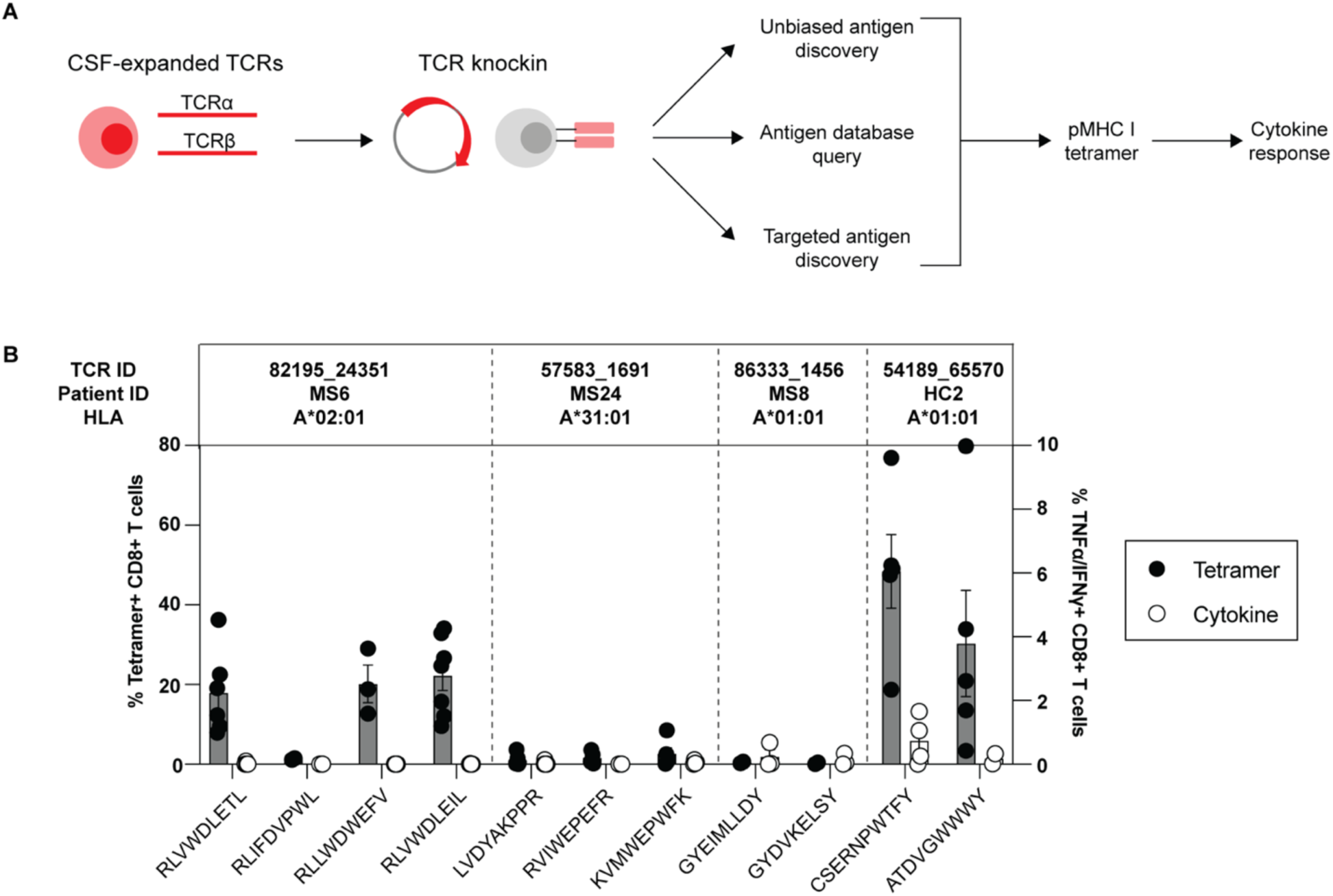
Antigen discovery of highly expanded CSF-enriched CD8+ T cells. Individual TCRαβ pairs were cloned into plasmids and expressed in primary human CD8+ T cells by non-viral CRISPR knockin. Candidate antigens for testing specificity were identified in three parallel strategies and were screened by pMHC tetramer binding and validated by cytokine production to cognate antigen (**A**). Candidate antigens for four TCRs identified by pMHC yeast display (unbiased antigen discovery) were tested for tetramer binding and cytokine reactivity (**B**). Each peptide was tested a minimum of two times using T cells from different donors for all tetramer and cytokine experiments.

Each TCR was expressed individually in primary human CD8+ T cells by non-viral CRISPR/Cas9-mediated TCR knockin as previously described^24^ (**Fig. S4A**). Candidate TCR-expressing CD8+ T cells were then probed for antigen specificity using pMHC I tetramers loaded with peptides identified from yeast display library screening. Three of the four tested TCRs demonstrated robust tetramer binding to most or all of the library-identified peptides (**Fig. 4B**, **Fig. S4B**). The ability of CD8+ T cells expressing these TCRs to respond functionally to the same antigens was tested by intracellular cytokine stimulation using antigen presenting cell (APC) lines expressing the relevant MHC I allele. Strikingly, only TCR clonotype 54189_65570 demonstrated cytokine production to peptide CSERNPWTFY, while none of the other TCR-expressing CD8+ T cells were functionally responsive to the respective yeast display-derived peptides (**Fig. 4B**). As nearly all the yeast display peptides identified by pMHC I tetramers were non-naturally occurring mimotopes, the analysis was extended to an array of foreign and human peptide homologs (**Table S9**). Varying degrees of tetramer binding were observed depending on the TCR tested, but none of the peptide homologs elicited cytokine responses above background (**Fig. S5A-B**). Thus, while the unbiased antigen discovery approach yielded novel mimotopes that identified several CSF-enriched CD8+ T cell clonotypes by pMHC I tetramer binding, none appeared to be genuine target antigens.

### Probing viral specificity of clonally expanded CD8+ T cells

The highly expanded CSF-enriched TCR clonotypes were queried against several public TCR databases, including VDJdb^25^, TCRex^26^, and TCRmatch^27^, as an additional TCR antigen discovery strategy (**Fig. 4A**). One CD8+ T cell clonotype (86333_1456) from participant MS8 demonstrated an exact match to both TCR α and β sequences with a well described the Epstein-Barr virus (EBV) epitope EBNA3A_193-201_ (FLRGRAYGL) (**Table 2**), which is restricted by HLA-B*08:01, an allele carried by this individual (**Table S1**). This identical clonotype was also moderately expanded in the CSF in an individual with Alzheimer’s disease^21^. A second CD8+ T cell clonotype (69317_24418) from participant MS6 was a near exact match to a TCR specific for BZLF1_54-64_ (EPLPQGQLTAY), another EBV epitope restricted by HLA-B*35:01, an allele also carried by this individual (**Table 2**).

**Table 2.**
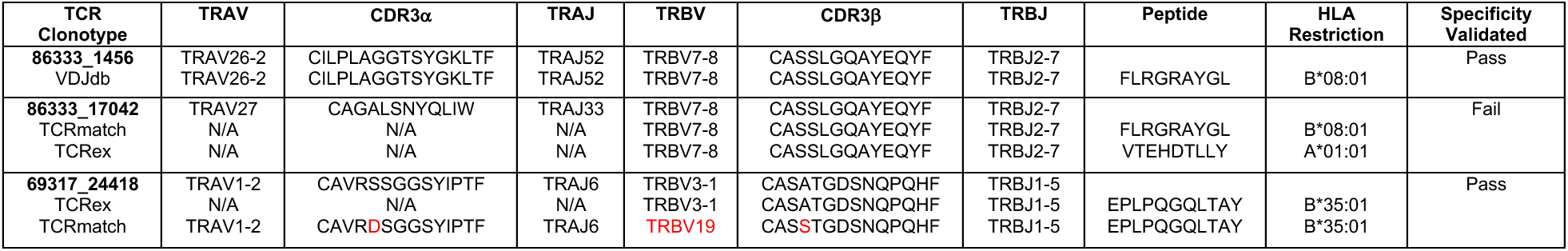
Results of CSF-enriched CD8+ TCRs queried against antigen-specific TCR sequences. The TCR clonotypes from the dataset (bold) that yielded at least a partial match to a TCR sequence from the indicated database is shown in each row. Differences in TCR genes and CDR3 sequences are indicated in red. N/A indicates TCR sequence not available. The peptide sequence and HLA restriction refer to the antigen linked to the database TCR sequence. Specificity validation of the indicated peptides was determined by pMHC tetramer binding and cytokine production to cognate antigen. Pass indicates successful validation and fail indicates the TCR clonotype ID was not specific for the indicated peptides.

These TCRs were expressed in primary human CD8+ T cells as above and their specificity was tested by pMHC I tetramer analysis. TCR 86333_1456 and TCR 69317_24418 showed robust tetramer staining to EBNA3A_193-201_:B*08:01 and BZLF1_54-64_:B*35:01, respectively (**Fig. 5A**). To confirm functional reactivity, human CD8+ T cells expressing each of these TCRs were stimulated with APC lines expressing cognate HLA and loaded with or without cognate EBV peptide. Each TCR demonstrated clear cytokine production to the relevant EBV peptide (**Fig. 5B-C**), confirming both CSF-expanded and enriched CD8+ T cell clonotypes are specific to EBV antigens.

**Fig. 5.**
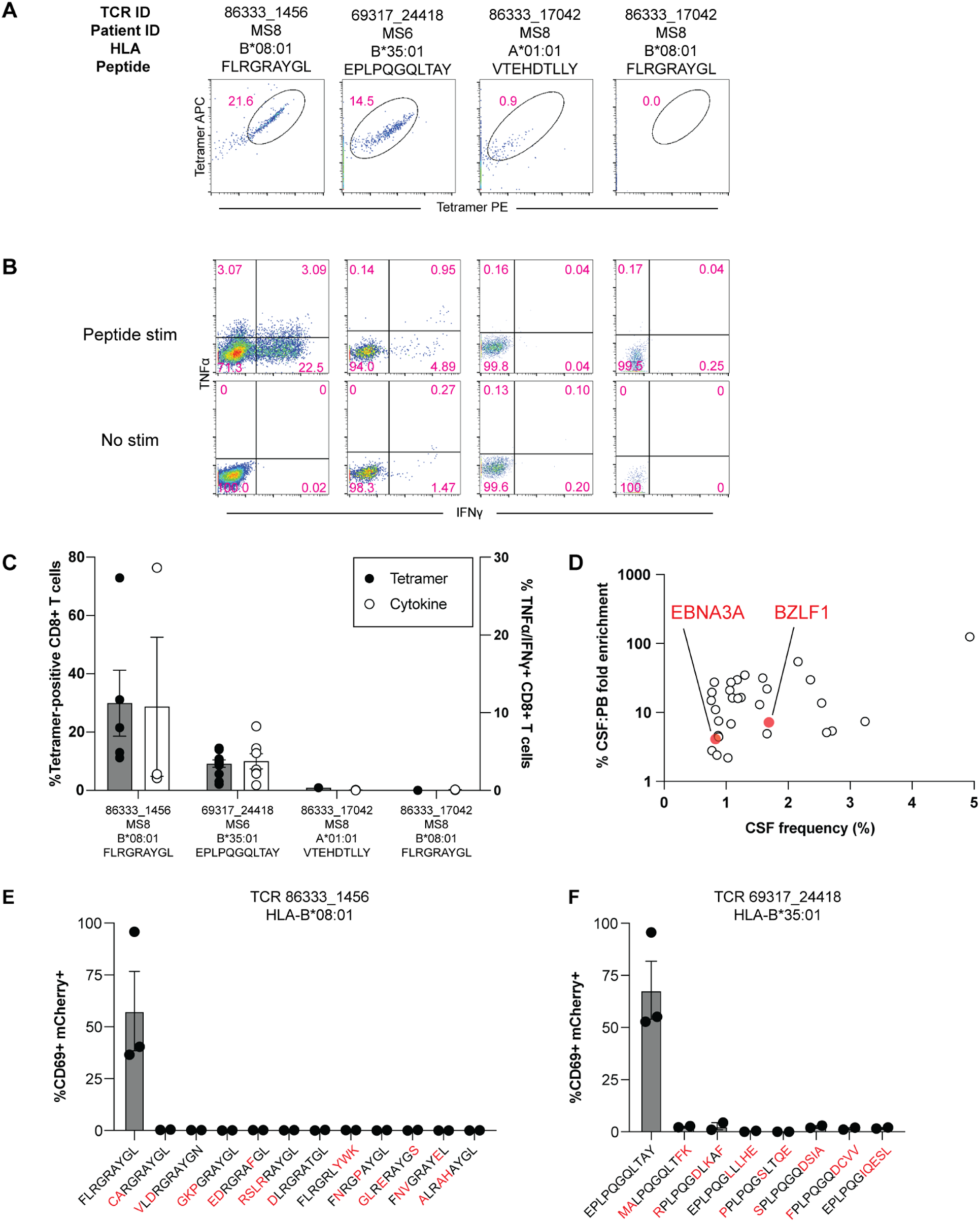
EBV specificity of highly expanded CSF-enriched CD8+ T cells. Representative flow cytometry analysis of tetramer binding (**A**) and cytokine production (**B**) for three MS patient TCRs with predicted reactivity to four different viral epitopes. Summary of tetramer binding and cytokine reactivity of each TCR (**C**) where cytokine reactivity reflects subtracted background from no stimulation control. FLRGRAYGL is EBV EBNA3A_193-201_ restricted by HLA-B*08:01, EPLPQGQLTAY is EBV BZLF1_54-64_ restricted by HLA-B*35:01, and VTEHDTLLY is CMV pp50_245-253_ restricted by HLA-A*01:01. The frequencies and degree of enrichment of the two EBV-specific clonotypes relative to all other highly enriched and expanded T cell clonotypes is shown (**D**). Summary of functional reactivity of Jurkat cells expressing the indicated TCR specific for EBV EBNA3A_193-201_:B*08:01 (**E**) or EBV BZLF1_54-64_:B*35:01 (**F**) to the indicated peptides. Responses reflect frequency of CD69/mCherry double-positive cells with no stimulation background control subtracted. Amino acid differences between cognate EBV peptides (left-most of each plot) and self-peptide homologs are indicated in red. Each peptide was tested in a minimum of two independent experiments.

In light of these findings, the possibility that additional CSF-enriched and expanded CD8+ T cells may be specific for viral antigens, in particular EBV, was further explored (SARS-CoV-2 peptides were not tested as all samples were collected prior to the COVID-19 pandemic). Nineteen TCRs were tested against panels of pMHC I tetramers loaded with previously identified immunodominant viral epitopes restricted by HLA matching that of the TCR donors. In total, 98 peptides restricted by 8 different MHC I alleles were screened **(Table S10**). Each TCR was screened with individual pMHC tetramers, except in the case of HLA-A*02:01 where tetramers were pooled in groups of 5 due to the large number of candidate peptides. Each TCR was tested against the indicated peptides a minimum of two times using two different T cell donors for TCR expression. No specific tetramer signal was observed for any of the TCRs to any of the peptides beyond the two EBV epitopes already identified for TCRs 86333_1456 and 69317_24418 (**Fig. S6**). Although TCR 86333_17042 from participant MS8 showed an identical match for a TCRβ sequence specific for EBV and cytomegalovirus (CMV) antigens with corresponding MHC I alleles (**Table 2**), it did not show any significant tetramer binding or cytokine reactivity to either viral antigen (**Fig. 5A-C**). These findings indicate that certain highly expanded CD8+ T cells in the CSF of MS patients are specific for EBV, but the specificities for most of the enriched T cell clonotypes remain unknown (**Fig. 5D**).

### Lack of self-antigen cross-reactivity of EBV-specific CD8+ T cells

To determine whether the two EBV-reactive CD8+ T cell clonotypes may be cross-reactive against self-antigens, the TCRs were screened against panels of self-peptides with partial sequence homology (**Table S11**). Using NFAT-mCherry-expressing Jurkat cells transfected with the CD8 co-receptor and TCR 86333_1456 or 69317_24418, high reactivity to the respective EBV peptides was confirmed (**Fig. 5E-F**). Strikingly, no significant reactivity was observed for any of the self-peptide homologs. While this does not entirely exclude the possibility for self-antigen cross-reactivity, it raises the possibility that the CSF enrichment of these clonally expanded CD8+ T cells may be driven by reactivity to EBV.

## DISCUSSION

CD8+ T cells are the dominant lymphocyte population in MS lesions^3,16^ where they are highly clonally expanded^5–8,12^, suggesting reactivity to hitherto unknown local antigens. Although several recent single-cell sequencing studies have explored gene expression changes and T cell clonal expansion in the CSF of MS patients^14–16^, significant questions remain regarding the identity of clonally expanded CD8+ T cells involved in MS and their antigen specificity. Our comprehensive transcriptional and clonal analysis identified CSF-infiltrating T cells with increased expression of genes associated with T cell activation, tissue resident memory (Trm), and CNS migration in treatment-naïve MS/CIS participants relative to controls, consistent with prior reports^14–17,19,28,29^. Due to the presence of clonally-expanded CD8+T cells in CSF in normal physiologic conditions and in CNS pathology^14,17,21^, identification of MS-specific CD8+ T cell clonal populations remains a challenge. Invoking a strategy previously used to identify disease-relevant T cells in inflammatory arthritis^20,30^ and cancer^31^, a subset of highly clonally expanded and CSF-enriched CD8+ T cells was identified that had the highest frequencies in MS/CIS participants. These CSF-enriched T cell clonotypes were widely characterized by a highly differentiated, antigen-experienced and cytotoxic phenotype with high CNS-trafficking potential. These gene signatures were very similar to the granzyme B and Trm markers enriched in CD8+ T cells in MS lesions^4,32,33^, strongly supporting the likelihood of these T cell clonotypes to be CNS-infiltrating.

Small networks of highly expanded CSF-enriched T cells with shared TCR sequence features to other less expanded clonotypes were found, which overwhelmingly occurred within the same individual. These findings suggest that distinct clonally expanded T cells may be contributory to MS pathology, unlike other recently described autoimmune conditions with preferential TCR usage^20^. Combined with the inherent technical challenges in T cell antigen discovery, these findings highlight the difficulties in identifying the antigen specificity of clonally expanded T cells in MS.

The majority of studies on candidate T cell autoantigens in MS have focused on CD4+ T cells^34–37^. By using three parallel antigen discovery strategies, our study provides significant new insight into the antigen specificity of CD8+ T cells in MS. Novel mimotopes to several MS-derived CD8+ TCRs were identified by pMHC yeast display, a powerful unbiased antigen screening tool. The majority of mimotopes were readily detectable by pMHC I tetramers loaded with the yeast display-derived peptides and naturally occurring peptide homologs, but only one elicited a measurable functional response. The reason for the discrepancy between pMHC tetramer binding and functional reactivity is unclear, but could be due to absence of catch bonds by certain high affinity TCR ligands^38^. Nonetheless, these candidate peptides provide an important framework for identifying TCRs with similar specificities in other individuals.

Two different CSF-expanded and enriched CD8+ T cell clonotypes specific for EBV antigens were identified. While EBV-specific CD8+ T cells have been previously reported in the CSF of MS and other neuro-inflammatory conditions^39–43^, the present study used paired TCRαβ analysis to unequivocally demonstrate EBV reactivity of highly enriched and clonally-expanded CD8+ T cell CSF populations in MS. These findings are particularly relevant in light of recent evidence that EBV infection is a prerequisite for the subsequent development of MS^44^. Interestingly, the EBNA3A:B*08:01-specific CD8+ TCR identified in one MS participant in this study was highly related to expanded CD8+ T cell clonotypes found in several patients with Alzheimer’s disease^21^. Our findings therefore provide further support that EBV may be related to multiple forms of CNS pathology.

The mechanism by which EBV is involved in MS pathogenesis remains unresolved. The fact that the EBV-specific CD8+ T cell clonotypes identified here were not only highly expanded but also preferentially enriched in the CSF of MS patients suggests these T cells may be responding to antigen within the CNS. EBV-specific B cells and CD4+ T cells in MS have been suggested to be cross-reactive to CNS autoantigens^45–47^ (i.e. molecular mimicry). However, we were unable to demonstrate cross-reactivity of the two EBV-specific CD8+ T cell clonotypes against partially homologous self-peptides, although this does not completely rule out such a mechanism. Alternatively, the findings of CD8+ T cells reactive against EBV late latent and lytic antigens are consistent with other reports^4,39,43^ and could reflect EBV reactivation within the CNS^48^.

The methodology of testing individual TCRs in primary human T cells by pMHC tetramer screening followed by validation with functional reactivity is highly rigorous and ensured only genuine positive results. This approach was particularly important in the case of a TCR that demonstrated an exact TCRβ match to another antigen-specific clonotype yet did not share the same specificity, highlighting the need to validate every TCRαβ individually. Antigen specificity should therefore be interpreted cautiously when based solely on partial TCR sequence matching.

This study was limited by a smaller population of control participants. Follow-up studies with larger numbers of well-matched MS and control participants are needed to more clearly identify disease-relevant T cell populations in MS. Although the transcriptional phenotyping analyses suggest a pro-inflammatory, cytotoxic phenotype of CSF-expanded CD8+ T cells, further *in vitro* and *in vivo* analyses are needed to determine whether these cells are pathogenic in MS. It is also important to acknowledge that despite the rigor of the antigen specificity testing, this approach was not exhaustive and was limited in the breadth of antigens that were tested. Given that various foreign and self-antigens are considered viable antigenic targets in MS, future studies will need to incorporate high throughput approaches to probe multiple target antigens simultaneously.

Elucidating the role of CD8+ T cells in MS requires 1) assessing their clonal repertoire in the CNS, 2) identifying their antigenic targets, and 3) determining their *in vivo* functions. This study provides significant progress towards all three aims by demonstrating a small population of predominantly CD8+ T cells that were highly expanded and enriched in the CSF of MS patients and strongly upregulated genes associated with antigen exposure, CNS migration, and cytotoxicity. The finding of EBV specificity of two of these CD8+ T cell clonal populations help to advance the understanding of MS pathogenesis and may pave the way toward potential antigen-directed therapies.

## MATERIALS AND METHODS

### Study cohort

MS/CIS and control participants were enrolled through the University of California San Francisco (UCSF) ORIGINS or Expression, Proteomics, Imaging, Clinical (EPIC) studies (https://epicstudy.ucsf.edu/). Healthy control and OND patients were enrolled in the biobanking study “Immunological Studies of Neurologic Subjects”. All enrolled MS and CIS participants were diagnosed according to the 2017 McDonald criteria^49^. Basic demographic and clinical information for all research participants is shown in **Table 1** and **Table S1**.

### Single-cell library preparation

CSF and blood were collected on the same day during clinical and research procedures in enrolled participants after informed consent. 20-30 mL of CSF was collected by lumbar puncture from each individual. Blood and CSF were processed immediately after collection in preparation for single-cell library preparation as previously described^18^. Unfractionated peripheral blood mononuclear cells (PBMCs) were isolated by CPT mononuclear cell preparation tubes (BD Biosciences) and resuspended in 2% fetal bovine serum. CSF was centrifuged at 400g for 15 minutes at 4°C, resuspended in ∼80 µL of supernatant, and counted. Single-cell sequencing libraries were prepared using 5’ scRNA-seq and 5’ T cell V(D)J scTCR-seq kits (10X Genomics).

### Single-cell sequencing analysis

Raw data for both scRNA-seq and scTCR-seq datasets were processed using Cell Ranger (v3.0.1 and v3.1.0, respectively) by 10X Genomics. The *cellranger count* and *cellranger vdj* commands were run with input Ensembl GRCh38.v93 and GRCh38.v94 references, for the scRNA-seq and scTCR-seq data respectively. All data were analyzed using a custom bioinformatics pipeline that included Seurat (v3.1.2 – v4.3.0), the Spliced Transcripts Alignment to a Reference (STAR) algorithm^50^ (v2.5.1), SingleR^51^ (v1.1.7), and DoubletFinder^52^ (v2.0.2). TCR V(D)J contig assemblies outputted from Cellranger were further annotated and analyzed using Immcantation (v3.1.0). TCR clonal families were identified using Change-O^53^ (v0.4.6), which generated clone IDs for both TCR alpha and beta chain assemblies.

### Quality control for single-cell data

Across both RNA-Seq and VDJ data, reads present in more than one sample that shared the same cell barcode and unique molecular identifier (UMI) were filtered using methods previously described^18^. The R package DropletUtils was used to filter out these reads in the RNA-Seq data and SingleCellVDJdecontamination (https://github.com/UCSF-Wilson-Lab/SingleCellVDJdecontamination) was used to apply the same methods to filter out these reads in the VDJ data.

All gene counts from scRNA-Seq data were combined using Seurat. Only genes present in two or more cells were included. Only cells containing transcripts for between 700 to 2,500 genes were included. The PercentageFeatureSet function was used to calculate the percentage of mitochondrial transcript expression for each cell. Cells which expressed at least 10% mitochondrial genes were omitted. Gene counts were normalized using the R package SCTransform^54^. The parameters **do.correct.umi** was set equal to TRUE and **var.to.regess** was set equal to nCount_RNA. All filtered cells were clustered using 20 principal components (PC) in Seurat. Clusters were formed using a shared nearest neighbor graph in combination with dimensional reduction using uniform manifold approximation and projection (UMAP)^55^, Doublet detection and removal were performed per sample using DoubletFinder^52^ with expected doublet rates set based upon the 10x Genomics reference manual. Cumulative sums were iteratively calculated for each PC to measure the percent variance accounted for with the data. To determine a reasonable number of PCs, a threshold of 90% variance was applied, which resulted in 12 PCs being inputted when re-clustering cells. Clusters of cells with a high expression of platelet markers, *PPBP* and *PF4*, or hemoglobin subunits (*HBB*, *HBA1*, *HBA2*) were omitted. Among the remaining cells, all V gene transcripts (*TRAV*, *TRBV*, *IGHV*, *IGKV* and *IGLV*) were removed and an additional round of re-clustering was performed with 9 PCs.

Assembled TCR contigs outputted from Cell Ranger were inputted into the Immcantation pipeline for a second round of alignments to the VDJ region using IgBLAST. Contigs containing fewer than three UMIs were omitted. Only contigs that aligned in frame (both the FUNCTIONAL and IN_FRAME output fields were TRUE) and across the constant region were retained. Cells in the TCR VDJ data were only kept if these contained one TCR beta chain and one TCR alpha chain. If cells had multiple chains, TCR alpha or beta, which passed these thresholds, the contig with the largest number of UMIs and or reads was kept.

### Cell type annotation and differential gene expression analysis

Cell types annotations were generated using methods previously described^18^. Cell types were defined by performing differential gene expression (DGE) analysis for each cluster. The normalized gene expression profile for each cluster was compared with the remaining cells using a Wilcoxon rank sum test using the FindAllMarkers function (min.pct = 0.1, logfc.threshold = 0.25, return.thresh = 0.01). The most up-regulated genes, with the highest positive average log fold change, were compared with a custom panel of canonical gene makers (**Table S12**) spanning several key immune cell types, including B cells, CD4+ T cells, CD8+ T cells, natural killer (NK) cells, classical monocytes, inflammatory monocytes, macrophage, plasmacytoid dendritic cells, and monocyte-derived dendritic cells. In addition to these manual cell type annotations, another set of cell types were determined using the automated cell type annotation tool SingleR, which used the combined Blueprint and ENCODE reference dataset for fine tuning predictions^51^.

A T cell subset was created by filtering for cells that overlap both RNA-Seq and TCR VDJ data. All clusters annotated as T cells had their annotations modified by CD8 gene expression. Among T cells, any cell with *CD8A* or *CD8B* expression was annotated as a CD8 T cell. The remaining T cells were then annotated as CD4 T cells. Differential gene expression (DGE) analyses were performed using the FindMarkers command in Seurat with the Wilcoxon test and the following parameters: P-adjusted value cutoff = 0.05 and logFC cutoff = 0.0.

### T cell immune repertoire analysis

TCR contigs outputted from Immcantation were clustered based on similarities between their T cell receptor variable region genes (*TRAV*, *TRBV*), T cell receptor joining region genes (*TRAJ*, *TRBJ*), and Complementary Determining Region 3 (CDR3) amino acid (AA) sequences. A TCR clone was defined as cells containing TCR alpha and beta chains which each contain identical V genes, J genes and CDR3 AA sequences. Cell counts were computed per clone ID, including separate cell counts for PBMC and CSF samples. Shannon’s entropy was calculated between CSF and PBMC in different disease groups using the alphaDiversity function in the R package alakazam. Specifically, the exponential of diversity scores (D) from the Shannon-Wiener index were extracted from the output of alphaDiversity by filtering for the diversity order (q = 1). Clonal expansion was defined as clones containing more than 1 cell. Among expanded clonal families of TCRs, CSF enrichment of highly expanded clones was determined by the ratio of the CSF to peripheral blood frequencies. Clones with a CSF frequency of ≥ 0.75% were annotated as CSF highly expanded. Clones that were expanded in CSF (i.e. more than singletons), but with frequencies less than highly expanded, were labeled as moderately expanded. Among expanded clonal families of TCRs, CSF enrichment of highly expanded clones was determined by the ratio of the CSF to peripheral blood frequencies. CSF highly expanded TCRs were inputted into GLIPH2 to generate glyph groups which indicated which TCRs were predicted to target the same epitope. These glyph groups were used to create a network using the R package igraph and graphically displayed using Cytoscape.

### HLA Genotyping

Sequencing for HLA was as previously described, adapted to include HLA class II^56^. For each sample, 100 ng of high-quality DNA was fragmented with the Library Preparation Enzymatic Fragmentation Kit 2.0 (Twist Bioscience). After fragmentation, DNA was repaired, and poly(A) tails were attached and ligated to Illumina-compatible dual index adapters with unique barcodes. After ligating, fragments were purified with 0.8x ratio AMPure XP magnetic beads (Beckman Coulter). Double size selection was performed (0.42x and 0.15x ratios), and libraries of approximately 800 bp were selected, at which point libraries were amplified and purified using magnetic beads. After fluorometric quantification, each sample was pooled (30 ng per sample) via ultrasonic acoustic energy. A Twist Target Enrichment Kit (Twist Bioscience) was then used to perform target capture on pooled samples. Sample volumes were then reduced using magnetic beads, and DNA libraries were bound to 1,394 biotinylated probes. Probes were designed specifically to target all exons, introns and regulatory regions of the classical HLA loci, including HLA-A, HLA-B, HLA-C, HLA-DPB1, HLA-DRB1 and HLA-DQB1. Then, streptavidin magnetic beads were used to capture fragments targeted by the probes. Captured fragments were then amplified and purified. Bioanalyzer (Agilent) was then used to analyze the enriched libraries. After evaluation, enriched libraries were sequenced using a paired-end 150-bp sequencing protocol on the NovaSeq platform (Illumina). After sequencing, HLA genotypes were predicted using HLA Explorer (Omixon).

### TCR cloning

The TCR sequences for each α and β gene pair were codon optimized and used to generate gene blocks (IDT) in which the TCRβ gene and TCRα were separated by a P2A sequence. Flanking homology arms were included to permit knockin into the human *TRAC* locus, as previously described^24^. The gene blocks were cloned into pUC19 plasmids by Gibson assembly and the correct sequence was verified by Sanger sequencing.

### Primary human T cell culture

Primary human CD8+ T cells were isolated from commercially purchased leukopaks (Vitalant or Stemcell) from unidentified healthy donors. PBMCs were isolated by Ficoll centrifugation and cryopreserved prior to each experiment. In all experiments T cells were cultured in RPMI containing with 10% fetal bovine serum, 2-mercaptoethanol, penicillin/streptomycin with L-glutamine, sodium pyruvate, MEM vitamin solution, and non-essential amino acids (all Fisher Scientific). TCR knockin was performed as previously described with minor changes^24^. In brief, CD8+ T cells were isolated from thawed PBMCs by negative selection (Miltenyi) and rested overnight with human IL-7 (5 ng/ml). CD8+ T cells were stimulated 1:1 with anti-human CD3/CD28 magnetic Dyna beads (Fisher Scientific), human IL-2 (20 ng/ml) and human IL-7 (5 ng/ml) and IL-15 (5 ng/ml) for 48 hours prior to T cell electroporation.

### Ribonucleoprotein (RNP) production for TCR knockin

gRNAs specific for the human *TRAC* locus were generated by incubating crRNA (AGAGTCTCTCAGCTGGTACA) 1:1 with tracrRNA (Dharmacon) for 30 minutes at 37 °C to yield a final concentration of 80 µM. Polyglutamic acid (PGA) was at 0.8 volume to the gRNA as previously described^57^. Cas9 (QB3 Macrolab) was added 1:1 with the gRNA and incubated for 15 minutes at 37 °C to yield a 20 µM RNP, which was immediately used for electroporation.

### TCR knockin of primary human T cells

48 hours after CD8+ T cell stimulation, Dyna beads were removed from the T cell culture using an EasySep separation magnet (StemCell). T cells were then spun at 200g for 9 minutes and resuspended in Lonza electroporation P3 buffer with supplement at 20 µl per 1 million T cells. 20 µl T cells were electroporated with 3.5 µl RNP and 1 µg of TCR-encoding plasmid DNA (1-2 µl) using a Lonza 4D Nucleofector 96-well electroporation system with pulse code EH115^58^. CD8+ T cells were immediately rescued by adding 80 µl warmed T cell media and incubating for 15 minutes in a 37 °C incubator. Cells were then split into fifths in 96-well round bottom plates and brought up to 200 µl with T cell media and 10 ng/ml IL-2. CD8+ T cells were expanded for a minimum of 96 hours before testing for pMHC tetramer binding. T cells were re-fed with a half volume of fresh media and IL-2 every 3-4 days.

### pMHC tetramer screening

Ultraviolet (UV) photolabile pMHC I monomers for HLA-A*01:01, HLA-A*A2:01, HLA-A*03:01, HLA-A*24:02, HLA-B*08:01, HLA-B*15:01, HLA-B*35:01, and HLA-B*44:02 were obtained from the NIH Tetramer Core. Custom peptide-loaded MHC I monomers were generated by UV-ligand exchange as previously described^59^. HLA-A*31:01 pMHC monomers (Easymers) were purchased from ImmunAware and loaded with custom peptides according to the supplier’s instructions. Tetramerization was carried out using streptavidin conjugated to fluorophores PE and APC (Life Technologies). CD8+ T cells were treated with 100 nM dasatinib (StemCell) for 30 min at 37 °C followed by staining with the appropriate tetramers (2-3 µg/mL) for 30 min at room temperature. All tetramers were used within 3-4 weeks of synthesis. Cells were washed in FACS buffer (1× DPBS without calcium or magnesium, 0.1% sodium azide, 2 mM EDTA, 1% FBS) and stained with anti-CD8 PECy7 (eBioscience; SK1), TCR BV421 (BioLegend; IP26), a PerCP/Cy5.5 dump antibody mixture containing anti-CD4 (BioLegend; RPA-T4), anti-CD14 (BioLegend; HCD14), anti-CD16 (BioLegend; B73.1), and anti-CD19 (BioLegend; HIB19), and Aqua506 viability dye (Life Technologies) for 30 minutes at 4°C. Cells were then washed and resuspended in FACS buffer and analayzed by flow cytometry (LSRFortessa). Only experiments where the FSC vs SSC gate contained at least 10% lymphocytes and where CD8+ T cells expressed less than 20% TCRs were used for analysis to ensure large number of T cells with high TCR knockout efficiency was achieved (**Figure S4**).

### pMHC yeast display selection

Yeast libraries were developed as previously described^23^. Yeast allele libraries were thawed in SDCAA (pH 5), passaged, induced in SGCAA (pH 5), and selected using biotinylated soluble TCR coupled to streptavidin-coated magnetic MACS beads (Miltenyi) as previously described^60^. Briefly, 2×10⁹ yeast cells from all four length libraries underwent negative selection with 250 μL of beads for 1 hour with rotation at 4°C in 5 mL of PBE (PBS + 0.5% bovine serum albumin + 1 mM EDTA). After passage through an LS column (Miltenyi) on a magnetic stand and three washes with 3 mL of PBE, the flow-through was incubated with rotation for 3 hours at 4°C with 250 μL of beads pre-incubated with 400 nM biotinylated TCR. The yeast were magnetically separated through an additional LS column, washed three times with 3 mL of PBE, and the elution was grown overnight in SDCAA (pH 5) following an SDCAA wash to remove residual PBE. Yeast were induced in SGCAA (pH 5) for 2-3 days before further selection, with subsequent selections using 50 μL of beads or TCR-coated beads in 500 μL of PBE.

### Deep sequencing of pMHC yeast libraries

DNA was isolated from 5-10×10⁷ yeast per selection using a Zymoprep II kit (Zymo Research). Unique barcodes and random 8mer sequences were added to the sequencing product by PCR and amplified for 25 cycles to allow for downstream demultiplexing and improved clustering. A subsequent PCR added Illumina chip primer sequences, resulting in products containing Illumina P5-Truseq read 1-(N8)-Barcode-pHLA-(N8)-Truseq read 2-IlluminaP7. The library was purified by double-sided SPRI bead isolation (Beckman Coulter), quantified using a KAPA library amplification kit (Illumina), and deep sequenced on an Illumina Miseq with a 2 x 150 V2 kit for low-diversity libraries.

### Generation of HLA-expressing APC lines

The genes for HLA-A*31:01, HLA-B*08:01 or HLA-B*35:01 were codon optimized and synthesized as gene blocks (IDT). The gene blocks were cloned into the pHR-CMV Lacz lentivirus vector by Gibson assembly and the sequence was verified by Sanger sequencing. The vector was expressed in K562 cells by lentiviral transduction and selected by puromycin.

### Cytokine assays

APCs for all cytokine assays were T2 cells expressing HLA-A*02:01, HLA-A*01:01 or HLA-A*03:01 or K562 cells expressing HLA-A*31:01, HLA-B*08:01 or HLA-B*35:01. APCs were pulsed with 10 µg/ml peptide or vehicle control overnight in serum-free media. CD8+ T cells (2 × 10^5^) were stimulated with peptide-loaded APCs (1 × 10^5^ per condition) for 6 hours in the presence of 1:500 GolgiStop (BD), 1:500 GolgiPlug (BD), and 1:200 CD28/CD49d (FastImmune; BD). Cells were washed with FACS buffer and stained with the cell surface antibodies as above (anti-CD8, anti-TCR, dump channel antibody mixture, and live/dead dye). Cells were washed, fixed, and stained with anti-human IFNγ Alexa 647 (BioLegend; 4S.B3) and anti-human TNFα Alexa 488 (BioLegend; Mab11) in permeabilization buffer (BD). Cells were then washed and collected on an LSRFortessa.

### Generation of TCR-expressing NFAT-mCherry Jurkat cells

Jurkat E6-1 T cells (ATCC TIB-152) were maintained in RPMI supplemented with L-glutamine and 10% FBS. Endogenous *TRAC* and *TRBC1* expression in Jurkat cells were knocked out with synthetic crRNAs designed using the Alt-R system (IDT) containing the following genomic targeting sequences: *TRBC1*: CGTAGAACTGGACTTGACAG; *TRAC*: CTTCAAGAGCAACAGTGCTG. crRNA was complexed with 1:1 tracrRNA (IDT) (0.2 nmol of each) followed by 0.1 nmol recombinant Cas9 protein (Macrolab, Berkeley CA). RNP were transduced into Jurkat T cells using the Amaxa P3 Primary Cell Nucleofector Kit (Lonza) using pulse code CK116. *TRAC* knockout was performed first and loss of surface TCRαβ expression was confirmed by flow cytometry. *TRAC* knockout cells underwent subsequent knockout of *TRBC1*, which had previously been shown to lead to loss of TCRαβ expression in a line with overexpressed TCRα. To track T cell receptor activation, a lentiviral vector was constructed that contained an NFAT transcriptional reporter NBV^61^ upstream of a minimal CMV reporter driving mCherry fluorescent maker expression, and constitutive expression of iRFP670 under a Pgk promoter provided a marker of transduction. Jurkat cells lacking endogenous TCRαβ expression were transduced with the vector and sorted for iRFP fluorescence and lack of mCherry background fluorescence. For TCRs corresponding to CD8+ T cells, an additional lentiviral vector encoding human CD8α was expressed in the Jurkat cells, and cells were sorted for uniform CD8α expression prior to TCR transduction. For TCR expression, lentiviral expression constructs were generated that encode a human Pgk promoter and the coding sequence of each specific TCR α chain with IRES-neomycin resistance gene or each TCR β chain with IRES-blasticidin resistance gene. Lentiviral particles were packaged in HEK293T cells following standard protocols and concentrated 10x using the Lenti-X Concentrator reagent (Takara). Viral particles were added to T cells at low multiplicity of infection, and expression was ensured by passaging cells for 5 days under antibiotic selection with 10 µg/ mL blasticidin (Gibco) and 1 mg/mL G418 (Teknova).

### TCR-expressing Jurkat cell assays

TCR-expressing Jurkats (1 × 10^5^) were stimulated for 24 hours with HLA allele transduced APCs (1 × 10^5^) loaded with 10 µg/ml peptide or vehicle control. Antigen-reactive CD8+ cells were identified by co-expression of NFAT-mCherry and anti-human CD69 PE (BioLegend; FN50).

### Statistical analyses

Shannon entropy results were compared using Brown-Forsythe and Welch ANOVA with multiple comparisons using Dunnett T3 corrections. Comparisons of CSF-enriched clonotypes T cell (e.g. CD8+ versus CD4+ T cells and MS/CIS vs HC/OND) were performed using unpaired two-tailed t-tests.

## Acknowledgments

University of California, San Francisco MS-EPIC Team: Anna Sindalovsky, Stacy Callier, Harkee Halait, Adam Santaniello, Adam Renschen, Simone Sacco MD, Nico Papinutto PhD, Ahmed Abdelhak MD, Ari Green MN, MCR, Joanne Guo MD, Sasha Gupta MD, Richard Cuneo MD, Jeffrey Gelfand MD MAS, Riley Bove MD MSc, Samuel Pleasure MD PhD, Roland G Henry PhD, Sergio Baranzini PhD, Jorge R Oksenberg PhD

We express gratitude to the individuals who agreed to participate as research subjects in this study. We thank the UCSF EPIC and Origins Study Teams for valuable aid in subject recruitment.

## Funding

Japan Society for the Promotion of Science (FH)

Japanese Society of Neurology (FH)

Mayer Foundation (JJS)

National Institutes of Health grant K08NS107619 (JJS)

National Institutes of Health grant R01AI158861 (JH)

National Institutes of Health grant R01AI169070 (JH)

National Institutes of Health grant R01NS092835 (SLH, MRW)

National Institutes of Health grant R35NS111644 (SLH, MRW)

National Institutes of Health grant R21AI142186 (MRW, SSZ)

National Multiple Sclerosis Society grant FAN-1608-25607 (RDS)

National Multiple Sclerosis Society grant FAN-1506-04555 (JJS)

National Multiple Sclerosis Society grant RG-2110-38434 (JJS)

Race to Erase MS (JJS)

Uehara Memorial Foundation (FH)

Valhalla Foundation (ORIGINS and EPIC cohorts)

Westridge Foundation (MRW)

## Author contributions

Conceptualization: BACC, SLH, SSZ, MRW, JJS

Participant recruitment: KK, TC, MH, RG

Methodology: KM, FH, RD, RDS, AR, JAH, MG, JJS

Investigation: KM, FH, RD, RDS, JG, LO, RL, AG, DAG, AR, ET, KK, JD, AC, FS, LF, TM, LLK, LS

Visualization: RD, JJS

Funding acquisition: JJS, SSZ, MRW, SLH, BACC

Supervision: JJS, SSZ, MRW

Writing – original draft: JJS

Writing – review & editing: BACC, SSZ, MRW

## Data and materials availability

All data are available in the main text or the supplementary materials.

