## Supplemental Figures 1-6 for "Antigen specificity of clonally-enriched CD8+ T cells in multiple sclerosis"

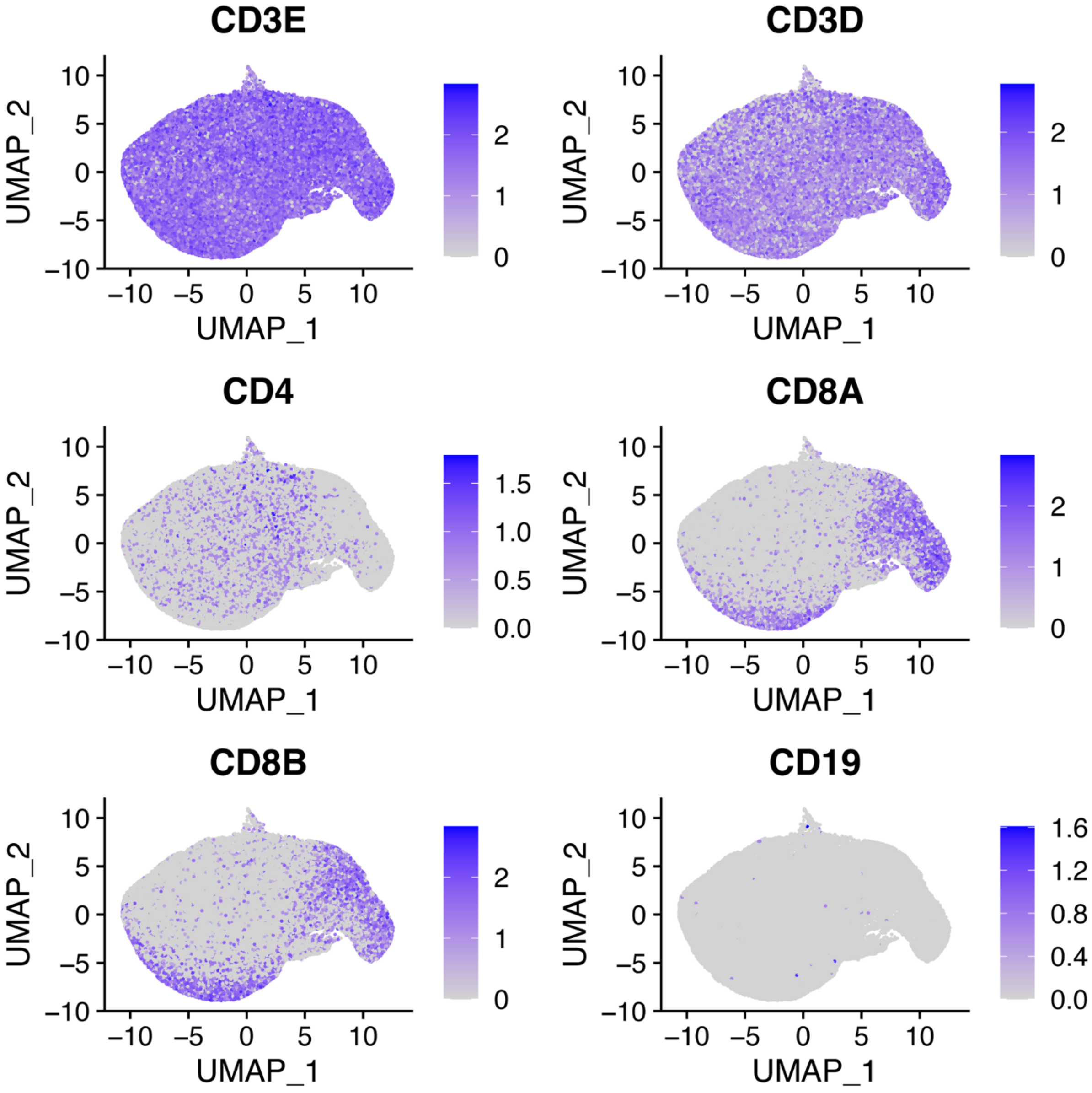


**Fig. S1. T cell gene expression analysis.** Expression for the indicated genes is shown for all T cells (blood and CSF combined) after merging scRNA-seq and scTCR-seq data.

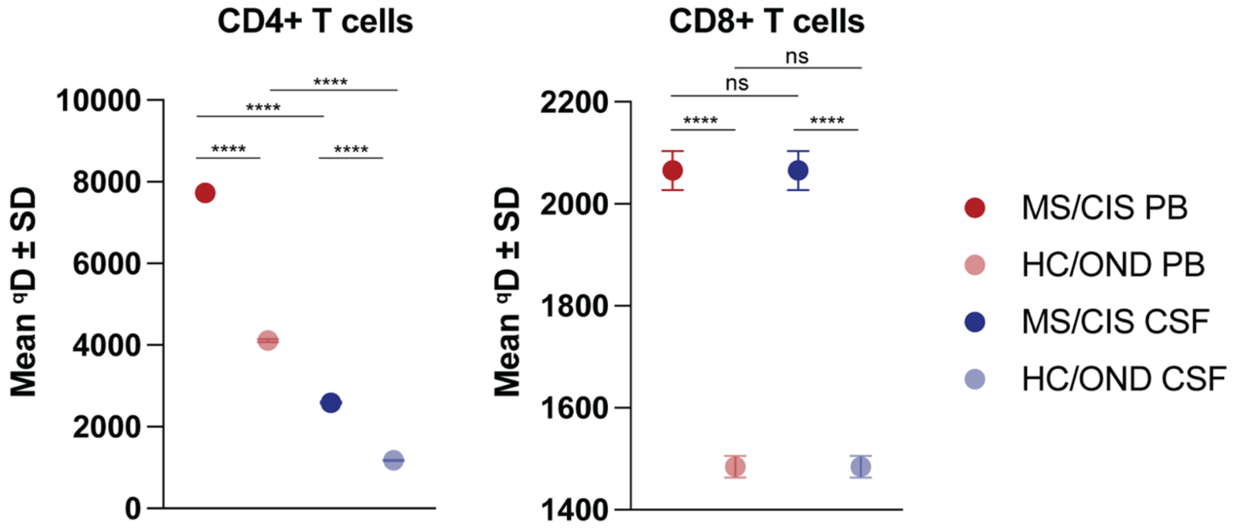


**Fig. S2. T cell diversity analysis.** Shannon entropy analysis (where y-axis indicates exponential of Shannon-Wiener index) is shown for CD8+ T cells in the peripheral blood (PB) and CSF by disease status. Abbreviations: MS = multiple sclerosis; CIS = clinically isolated syndrome; HC = healthy control; OND = other neuroinflammatory disease. ****$p < 0.0001$.

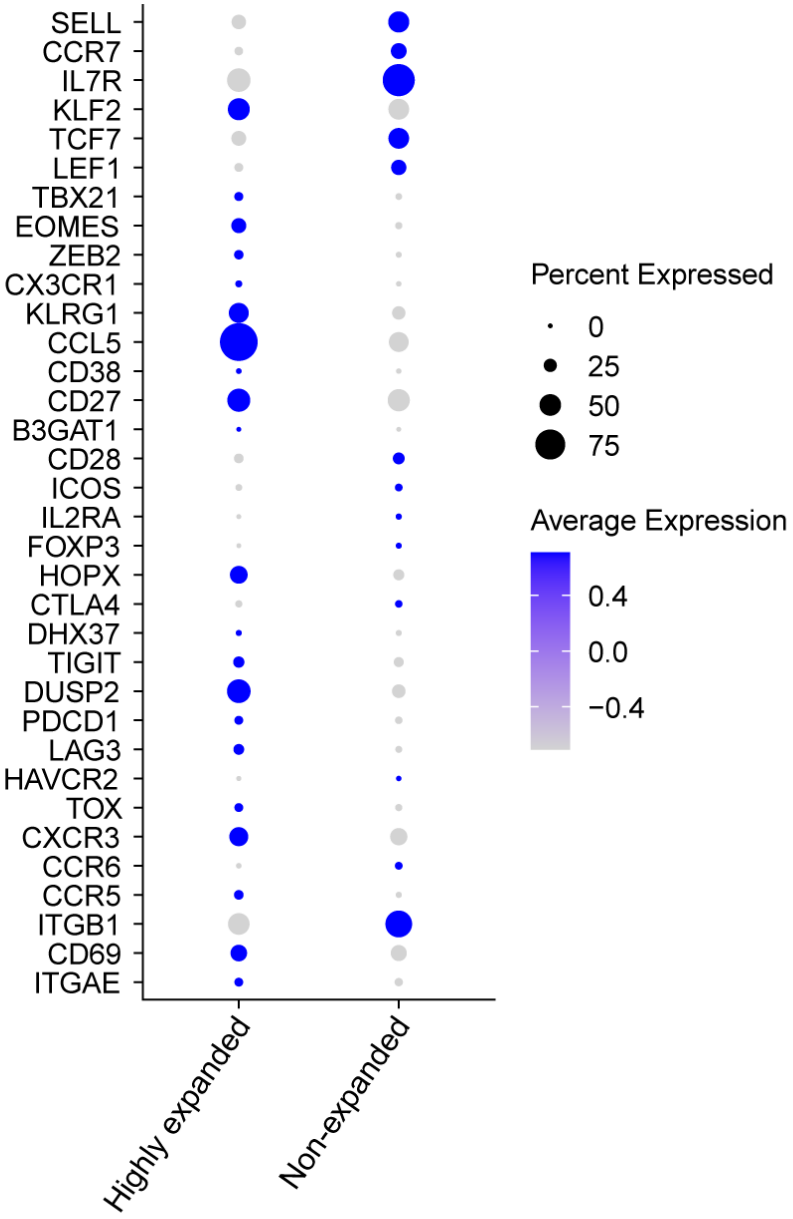


**Fig. S3. Gene expression analysis of CSF T cells by expansion status.** Expression levels and percent expression of the indicated genes is shown for highly expanded CSF T cells versus non-expanded T cell clonotypes.

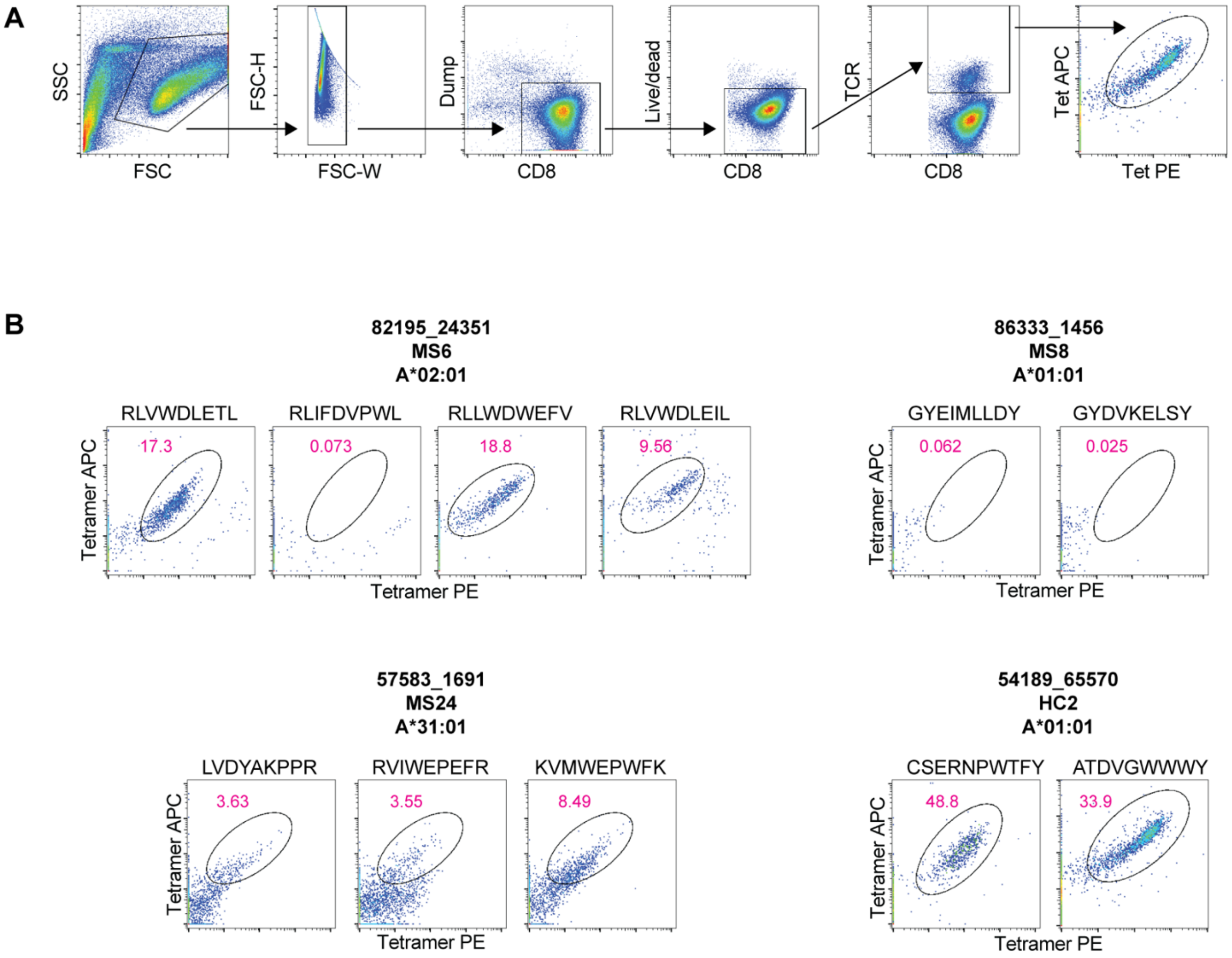


**Fig. S4. Tetramer screening of antigens identified by pMHC yeast display.** Representative flow cytometry analysis of pMHC tetramer stained CD8+ T cells following TCR knockin (**A**). Four patient-derived TCRs were tested for tetramer binding to the indicated peptides (**B**).

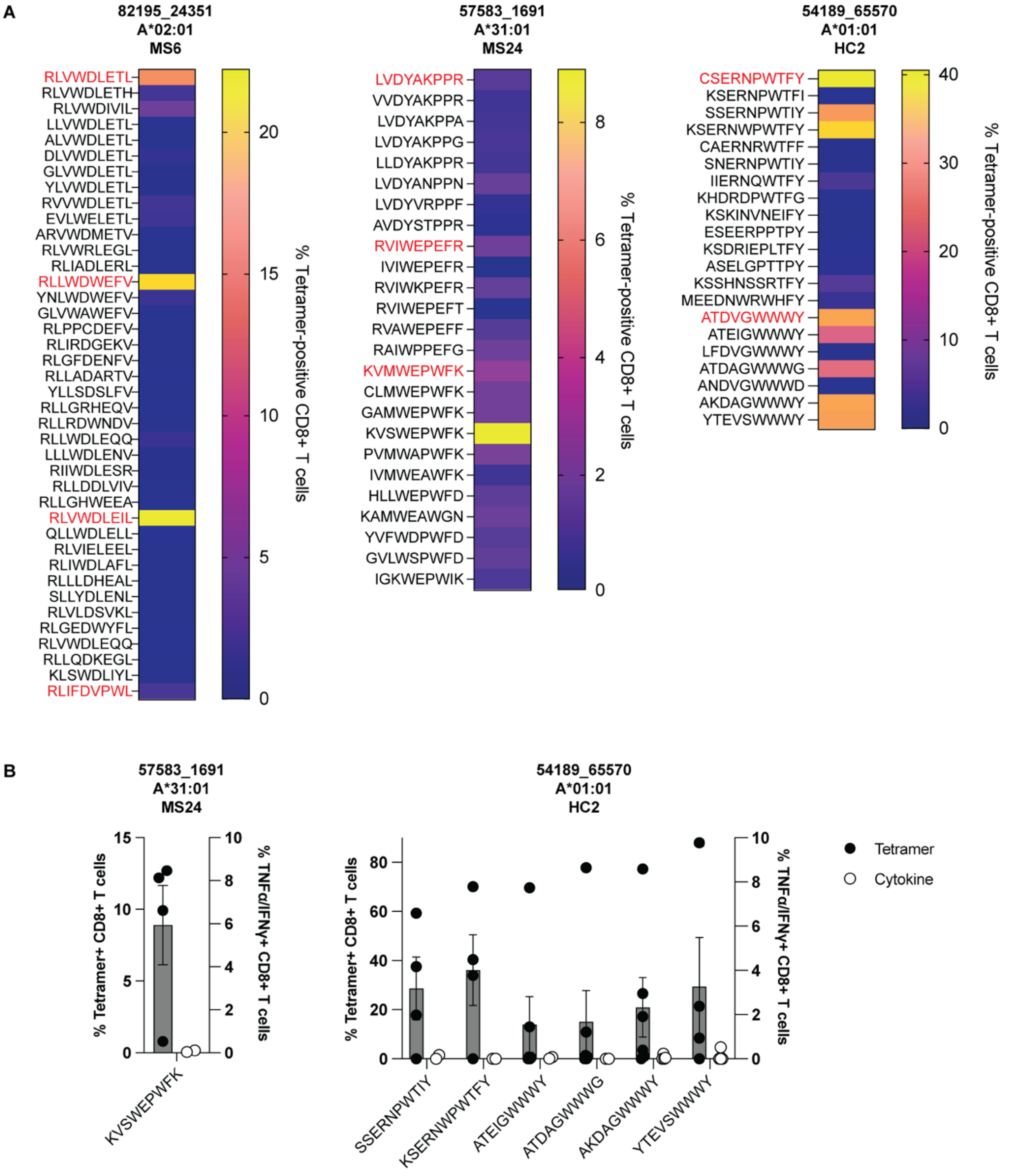


**Fig. S5. Validation results of peptide homologs to yeast display mimotopes.**

Summary tetramer binding analysis of peptide homologs for the three TCRs that yielded yeast display-derived mimotopes (red) (**A**). Summary of tetramer binding and cytokine reactivity of the indicated peptides to two different TCRs (**B**). Cytokine reactivity reflects subtracted background from no stimulation control. Each peptide was tested a minimum of two times using T cells from different donors for all tetramer and cytokine experiments.

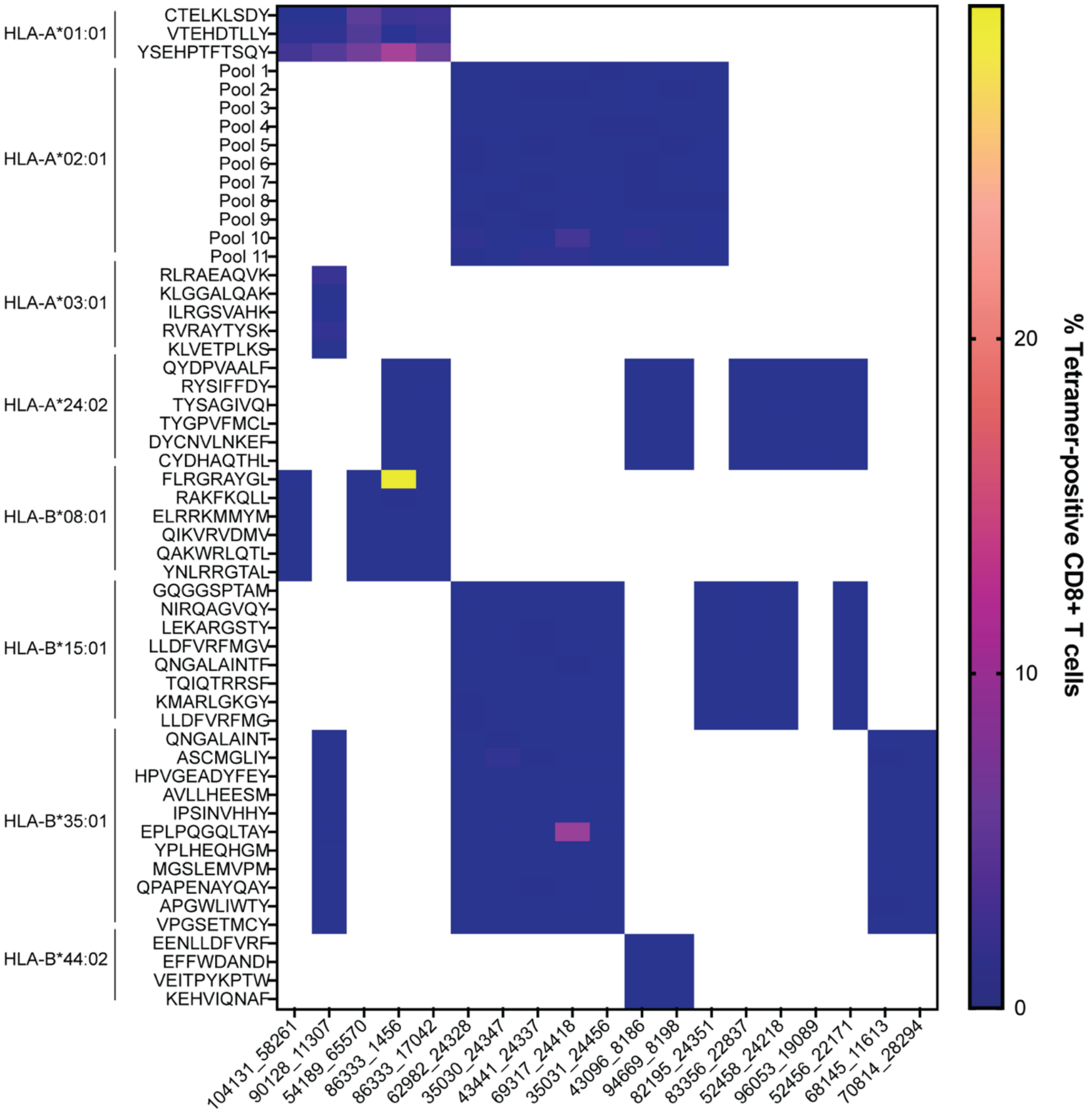


**Fig. S6. Summary of viral antigen specificity of highly expanded CSF-enriched CD8+ T cells.** The summary of all pMHC tetramer screening for 98 viral peptides for 19 patient-derived TCRs is shown in the heatmap. All viral peptides were tested individually except in the case of the peptides for HLA-A*02:01 where tetramers were tested in pools of 5. Each peptide/pool was tested a minimum of two times in using different T cells from different donors.
